# Born on Mars: Multigenerational phenotypic change in *Caenorhabditis elegans* under Martian analog gravity and hypomagnetic fields

**DOI:** 10.1101/2024.10.18.619154

**Authors:** A Akinosho, Z Benefield, X Dai, W Stein, AG Vidal-Gadea

## Abstract

Life on Mars will require organisms to endure sustained exposure to reduced gravity and near-absent planetary magnetism, yet little is known about how these environmental factors may combine to potentially affect biology across generations. Here, we investigated the transgenerational effects of Martian gravity and magnetism on neuromuscular, sensory, and morphological function using *Caenorhabditis elegans* as a model. Animals were reared continuously under ground-based simulated Martian conditions across six generations and assessed using high-throughput behavioral and morphometric assays. Animals remained viable but acquired progressive transgenerational impairments across multiple functional domains. Swimming frequency showed immediate and severe deficits at all generations tested (Cohen’s d = 2.6-4.2), while chemotaxis deficits emerged more gradually, becoming significant by Generation 4. Morphological changes followed non-monotonic trajectories, with transient compensatory growth at Generation 4 followed by increased developmental variability at Generation 6. These differential patterns of impairment across neuromuscular, sensory, and developmental systems reveal domain-specific vulnerabilities to sustained Martian conditions. Critically, by Generation 6, Mars lineages exhibited a three-to-eight-fold increase in phenotypic variability, consistent with a loss of developmental canalization that could pose a more serious challenge for sustained colonization than simple mean fitness reductions.

## Introduction

The planned human exploration of Mars will expose humans and other organisms to low-gravity and hypomagnetic environments. Mars has only ∼38% of Earth’s gravity and a negligible magnetic field with only localized crustal magnetism (Nimmo and Tanaka, 2005). Gravity is a pervasive physical cue that has guided biological evolution on Earth and plays a key role in many biological functions. Likewise, organisms have been continuously exposed to Earth’s geomagnetic field, and it plays subtle but significant roles in physiological processes, with hypomagnetic field exposure linked to oxidative stress, altered redox homeostasis, and behavioral changes in animal models (Tian et al., 2022, 2024; Zhang and Tian, 2020). Historical work established early interest in biomagnetic effects (Dubrov, 1978), and recent studies have clarified molecular mechanisms. Removing or altering gravity and magnetic field individually can have profound effects on living systems. For example, in microgravity, astronauts and other mammals experience muscle atrophy, bone density loss, and immune dysfunction (Grimm et al., 2016; Moosavi et al., 2021). Animal studies have identified hypomagnetic fields (HMF) as potential stressors that are linked to oxidative stress and cellular dysregulation (Tian et al., 2022, 2024). Understanding how life evolved on earth responds to reduced gravity and magnetic fields is thus not only critical for the medical treatment and multigenerational colonization by future astronauts but also offers insight into fundamental biology under extreme conditions.

Research across diverse organisms highlights the importance of gravity and magnetic cues across many animal species. Jellyfish reared in space, for instance, develop structural gravity-sensing organs (statoliths) that appear normal in microgravity, yet upon return to Earth these space-raised jellyfish show abnormal swimming and orientation behaviors and fail to navigate gravity properly (Spangenberg et al., 1994). This suggests that normal gravity is required for proper vestibular development and may represent a cautionary analog for human infants in space. Fruit flies (*Drosophila melanogaster*) flown on the Space Shuttle exhibit suppressed innate immune function, mirroring the immune system weakening seen in astronauts (Marcu et al., 2011). Space-flown mice and other mammals likewise show conserved responses such as muscle degeneration and metabolic shifts, underscoring cross-species commonalities (Gambara et al., 2017). Conversely, some organisms show partial compensatory responses: For example, some plants and small animals adjust their growth or behaviors when gravity is altered, although not without trade-offs (Cialdai et al., 2023; Maffei et al., 2024).

The nematode worm *Caenorhabditis elegans* has emerged as a powerful model for space biology because of its genetic tractability, physiological parallels with higher animals (Ishioka and Higashibata, 2022), and ability to weather spaceflight conditions. Decades of studies demonstrate that *C. elegans* can survive and reproduce in spaceflight and simulated microgravity, with microgravity and radiation eliciting molecular and physiological changes (muscle atrophy, metabolic reprogramming, etc.) that parallel those seen in many other animals. Worms flown on the Space Shuttle and International Space Station (ISS) generally complete their life cycle normally even in microgravity. For example, key processes like development and germline apoptosis occur at normal rates during short-term spaceflight (Honda et al., 2014). Early space experiments (e.g., shuttle mission STS-107) showed that worms not only lived through the flight but also produced offspring, indicating baseline viability in microgravity (Campbell, 2004). Notably, Selch et al. (2008) examined gene expression in space-flown worms and found that while developmental timing appeared normal, there were significant transcriptional changes in muscle development and metabolism, including a shift consistent with insulin/IGF-1 and TGF-β signaling alterations (Selch et al., 2008). Szewczyk et al. (2005) reported that worms exposed to spaceflight showed no lethal phenotypes and maintained normal behavior despite exposure to space radiation, demonstrating remarkable stress tolerance. Together, these studies established *C. elegans* as a robust organism for studying non-lethal, generational effects of the space environment. They also indicated that microgravity induces subtle stress responses even if outward development and survival remained grossly intact. With respect to lasting colonization of planets, studying multigenerational exposure becomes and important issue because some effects may accumulate or only emerge over longer timescales, for example through epigenetic changes or transgenerational adaptation. *C. elegans* is particularly well suited for this kind of work because it can transmit environmentally induced behavioral responses across generations through small RNA mediated signaling pathways. Moore et al. (2019) showed that learned pathogen avoidance can be inherited through the F4 generation via Piwi/PRG-1 argonaute and TGF-β pathways. This transgenerational effect was recently independently validated through the F2 generation by Akinosho et al. (2025), confirming the robustness of small RNA-mediated behavioral inheritance in *C. elegans*.

While characterizing the impact of individual environmental factors is a necessity, understanding the effects of combined environmental stressors for long-term space colonization is even more critical because biological systems often exhibit non-linear responses when multiple factors interact. More detailed studies in *C. elegans* showed that microgravity alone affects muscle and metabolic pathways, including reduced body length, muscle atrophy, and altered expression of cytoskeletal and mitochondrial genes (Selch et al., 2008; Çelen et al., 2023). Like many other animals, *C. elegans* uses the magnetic field to navigate its environment (Vidal-Gadea et al., 2015), and while the effects of changes to the magnetic field changes are not understood, studies in mammals have demonstrated that hypomagnetic fields independently disrupt redox homeostasis and neural function (Tian et al., 2022, 2024). The combined impact of Mars level gravity and magnetic field over multiple generations, however, remains unknown in any animal model. Surface gravity and the planetary magnetic environment are global features that cannot be directly engineered during colonization of extraterrestrial planets, at least not at the scale of whole organisms. Rather than dissecting the independent contributions of each field, here we therefore take a planet centric approach and ask whether the combined Martian physical environment (effective surface gravity plus magnetic field strength and geometry) is sufficient to drive cumulative heritable changes in an Earth-evolved animal.

We hypothesized that prolonged exposure to combined Martian gravity and magnetism would induce transgenerational physiological changes, but that different biological systems might exhibit distinct temporal patterns of impairment based on their energetic demands and buffering capacities. We predicted that high-energy neuromuscular functions (such as swimming) would show immediate vulnerability, while sensory-neural systems (such as chemotaxis) might exhibit delayed cumulative effects, and developmental programs might display complex adaptive responses. We tested these hypotheses by measuring morphological parameters, locomotor performance under varying mechanical demands (e.g., crawling versus swimming), and chemosensory function in Earth and Mars conditions across six generations. Each condition was propagated through independent biological lineages tracked longitudinally to enable detection of progressive transgenerational effects.

Our approach uniquely combines clinostat-based gravity simulation with magnetic field cancellation, enabling a novel examination of how Martian gravity and magnetism jointly shape biological function. We describe our simulation system, report the morphological, behavioral, and sensory outcomes observed, and discuss their relevance for long-term space habitation and biological adaptation beyond Earth.

## Methods

### Animal strains

In this study we used (N2) wild-type *C. elegans* nematodes obtained from the *Caenorhabditis* Genetic Center. Animals were raised on OP50 *E. coli* bacteria at 20±2°C following established protocols (Brenner, 1974).

### Animal husbandry

To study animals across six generations, approximately 3-4 day 1 adult worms were allowed to lay eggs in plates containing nematode growth media (NGM) agar seeded with a lawn of OP50 *E. coli* for one hour. Plates with freshly laid eggs were placed inside a PVC receptacle at the center of our experimental setup (see below). After 3.5 days plates were removed from the experimental setup and day-1 adults were once again allowed to lay eggs for one hour in new NGM plates before these were replaced in the setup once more to continue the transgenerational experiment. Following the 1-hour egg laying protocol, adult day-1 worms were used for the behavioral and physiological experiments. For each condition (Mars or Earth) four individual NGM plates with worms were propagated independently for six generations.

### Experimental setup

To manipulate the effective magnetic and gravitational field experienced by developing nematodes we constructed two identical experimental setups (Figure 1). Each setup consisted of a clinostat (BIO244, Eisco Labs) used to manipulate the experienced gravitational field, plus a 20 cm^3^ four-coil Merritt magnetic cage used to manipulate the magnetic field experienced by the animals. We used a four-coil Merritt system as these have been shown to produce the greatest homogeneous magnetic field volume to cage dimensions ratio (Merritt et al., 1983; Cao et al., 2018). The animals sat at the center of this cage surrounded by a grounded Faraday cage comprised of copper fabric (mesh <1mm) to cancel out the electric field generated by the magnetic coil system. Because our aim was to model the net physical environment at the Martian surface rather than to isolate the effects of single parameters, all Mars treated lineages experienced the combined gravity and magnetic field configuration; we did not include conditions with altered gravity alone or altered magnetic field alone.

**Figure 1.**
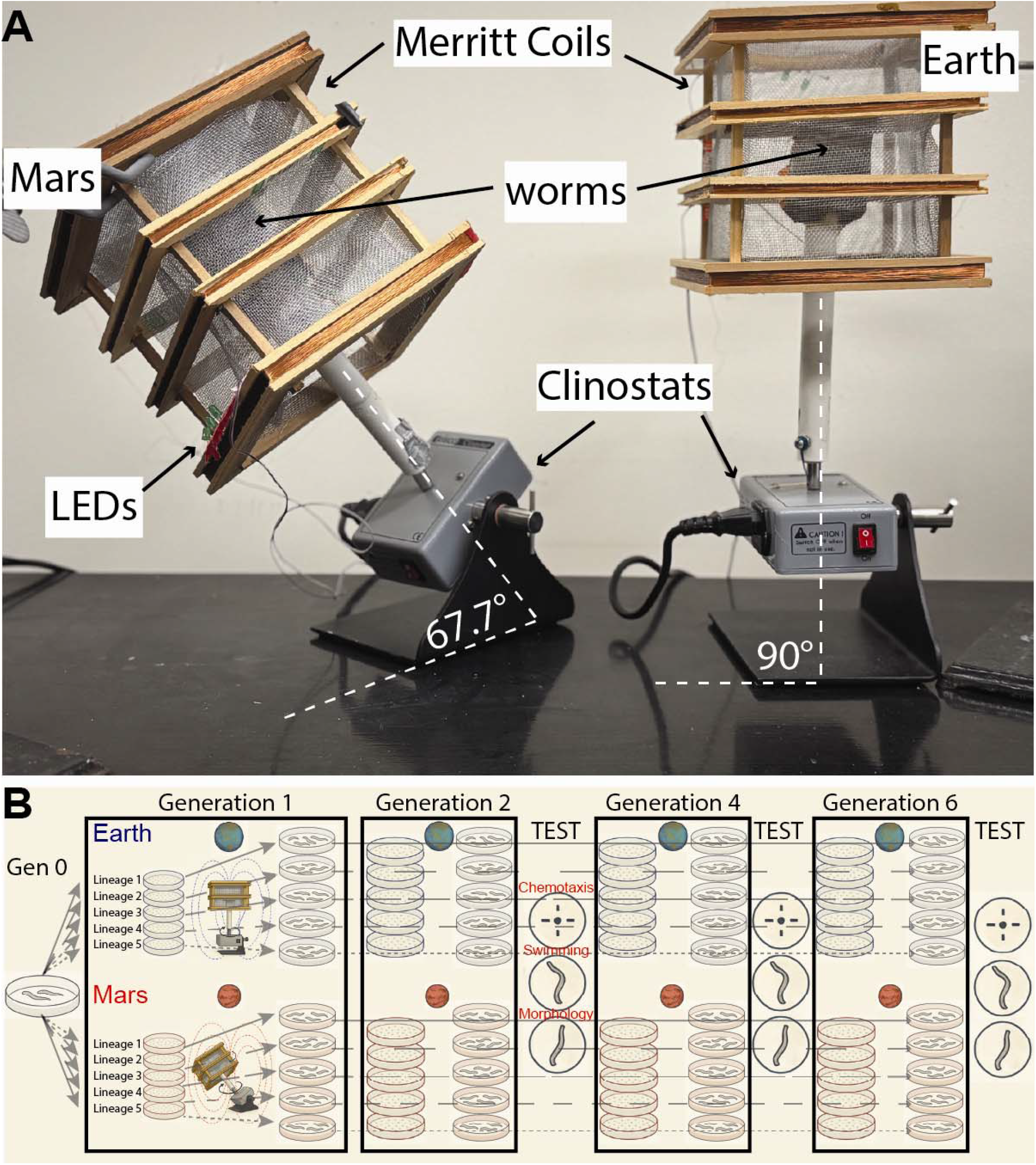
Simulation of Mars and Earth’s magnetic and gravitational fields. **A)** Two clinostats were positioned at different angles to either partially cancel the Earth’s effective gravitational pull (Mars: 67.7° from horizontal) or as a control (Earth: 90° from horizontal). Two identical 20cm^3^ 4-coil Merritt coil magnetic cages with internal (grounded) Faraday insulation. Both coils were powered by the same amplifier and current. Each coil system was angled so that the magnetic field produced at its center combined with the local Earth field to produce a net field mimicking Mars’ magnetic field (<6 mG), or Earth’s magnetic field (650 mG). In each system 3cm agar plates containing *C. elegans* eggs were placed and allowed to grow to adulthood at 20°C feeding on OP50 E. coli (3.5 days). The magnetic field within the systems was tested at the start and end of each experiment and LED indicator was used throughout to monitor the system while in operation. **B)** Experimental design. Synchronized populations of *C. elegans* were maintained under simulated Earth or Mars conditions continuously for six generations. At Generations 2, 4, and 6, day 1 adults were removed for behavioral and morphological assays. Each condition was propagated through 4-5 independent biological lineages (assay plates) tracked across all generations, creating a repeated measures design for statistical analysis.

In each of the two identical systems, the clinostat inclination and the Merritt cage orientation could be controlled independently. Both Merritt Cages were powered by a single DP50V5A constant voltage constant current programmable control power supply which delivered the same current to each of the two Merritt cages. The effective magnetic field at the center of each magnetic cage was measured using a DC milliGauss (mG) meter (AlphaLab Inc). The magnetic cages were aligned so that the magnetic vector they generated summed with the earth’s magnetic vector in the lab to produce a net magnetic field of <6±5 mG (Martian conditions), or 650±10 mG (Earth conditions). Magnetic conditions within the cages were measured before the start and at the end of each generational experiment. The electric field generated by the cages was removed by grounded Faraday cages inside the Merritt systems.

To ensure continuous and reliable operation of the magnetic coil system across the multi-generational experiment, LED indicators were integrated into each Merritt cage to provide real-time visual confirmation of power delivery throughout the 3.5-day developmental period. This modification was implemented after an initial pilot experiment was halted at Generation 4 due to concerns about potential equipment malfunction when Mars-reared animals unexpectedly exhibited larger body dimensions than Earth controls. However, with the enhanced monitoring system confirming continuous coil operation, a second complete experiment reproduced the Generation 4 morphological pattern, validating it as a genuine biological response rather than technical artifact.

To alter the effective gravitational pull experienced by the nematodes we calculated the angle α at which each clinostat needed to be positioned to model each net gravitational pull. Martian gravity is approximately 0.38g (38% that of the Earth’s). α for Martian gravity was calculated using Newtonian mechanics: sin(α) = (Martian gravity)/(Earth’s gravity). α was thus = 22.3°. For Earth gravity conditions, the clinostat was angled at 90° to the ground. Importantly, the clinostats rotated at a speed of 1 revolution per hour which implies that only processes occurring at and above these time scales could be impacted by this manipulation.

### Filming

#### Swimming

Plates containing Day 1 adult *C. elegans* were flooded with 2 ml of liquid NGM solution. Animals were then placed in a stereoscope microscope (Olympus CX12) and filmed for 30 seconds at 30 fps using a Pointgray USBC video camera. TIFF sequences were saved for later analysis. *Crawling*. Day 1 adults were transferred to NGM plates with no bacteria. A Basler acA4024-29um infrared camera was used to generate 17 sec 30 fps films of nematodes crawling.

### Chemotaxis assay

Chemotaxis was assessed using standard procedures (Queirós et al., 2021). Day one *C. elegans* were placed at the center of 10cm chemotaxis agar plates with 1 ml drop of 1×10^−3^M diacetyl in liquid Nematode Growth Media (NGM) as an attractant source at one side, and a 1 ml control drop of NGM at the opposite side. A 1 ml of the paralytic NaN_3_ was also added at each target point to paralyze and count animals following their migration. After 30 minutes at 22 ± 1°C, the number of worms on each side (attractant and control) was quantified. The chemotaxis index defined as C.I. = (A - C)/(A + C) (where A = number of animals at attractant site, and C = number of animals at control site) was calculated to evaluate the response. All assays were performed with five independent biological lineages per condition, with each lineage assayed at Generations 2, 4, and 6, following established protocols (Margie et al., 2013).

### Swimming frequency

Image sequences were manually analyzed using ImageJ. Swimming frequency was quantified from five independent biological lineages (assay plates) per condition at each generation. For each lineage, 7 individual worms were manually analyzed using ImageJ, yielding 35 total worms per condition per generation. The maximal number of uninterrupted swimming cycles produced by each animal was divided by the time elapsed during the interval to obtain swimming frequency.

### Tierpsy kinematic analysis

We used an automated behavioral analysis system to quantify morphology and locomotion of crawling worms. Day 1 adult worms were transferred to NGM plates without bacteria and filmed using a Basler acA4024-29 μm infrared camera mounted above the plate. A collimated LED light source provided high-contrast illumination from below, enabling clear visualization of worm body contours. Each test plate consisted of 15 Day 1 adult worms derived from an individual culture plate and was filmed for 20 seconds at 30 fps.

Video recordings were analyzed using Tierpsy Tracker software (Javer et al., 2018), which automatically detects and segments individual animals, extracts body contours, and generates skeletonized representations of each worm. Tierpsy quantifies over 4,600 metrics from each animal, including morphological parameters (body length, midbody width, body area at 10th, 50th, and 90th percentiles) and kinematic features (speed, angular velocity, body curvature, etc.). The software outputs the average result for each metric for the population sampled. Four replicate plates were analyzed per condition per generation, providing biological replicates for statistical analysis.

### Statistical analysis

All statistical analyses were performed using Python 3.12 with SciPy statistical packages. Complete statistical analysis code is provided in Supplementary Software 1, with all calculations independently verified in Supplementary Data 2. Our experimental design involved propagating four to five independent biological lineages (assay plates) per condition across generations, with each lineage measured at Generations 2, 4, and 6. This repeated measures structure creates non-independence among measurements from the same lineage across time. Because our primary interest was in trajectory differences between conditions at each time point, and because our sample size per generation was limited, we treated lineage means as independent observations within each generation and did not explicitly model within lineage correlations across time. Unless otherwise noted, data are presented as mean ± standard error of the mean (SEM).

Given the effect sizes observed across our measured traits, our sample size of n = 5 lineages per condition yields statistical power that varies by trait and reflects the relative biological sensitivity of different systems to Martian conditions. Swimming frequency, which showed the most severe and consistent impairment, has excellent statistical power (approximately 95 - 100% across all generations, Cohen’s d = 2.6 - 4.2), ensuring reliable detection of neuromuscular deficits. Chemotaxis shows progressive effects, with power ranging from about 26% at Generation 2 (where no difference was detected), to intermediate power at Generation 4, and to essentially 100% at Generation 6, appropriately reflecting the delayed onset of large sensory impairments. Morphological traits show moderate power (about 61 - 79%) for the comparisons that reached significance (body length at Generations 2 and 6; body area and midbody width at Generation 4), indicating that the detected effects represent robust biological changes despite modest sample sizes. Importantly, all six non significant comparisons had low statistical power (all below 50%), so these outcomes cannot be interpreted as evidence of equivalence between Earth and Mars conditions. Rather, they indicate that our design was tuned to detect only large effects, and any undetected differences are likely to be smaller than the very large effects we observed for swimming and late generation chemotaxis. The magnitude and consistency of detected effects across multiple independent traits and generations argue against Type I error from isolated significant comparisons. While a formal mixed effects model explicitly accounting for within lineage correlations across time would in principle provide a more complete statistical treatment, the present design with five lineages per condition does not provide enough replication for a stable implementation of such models, and given the very large and internally consistent effect sizes observed, such an approach is unlikely to change our biological conclusions.

We compared Earth versus Mars worms at each generation (Generations 2, 4 and 6) using planned contrasts implemented as two tailed t-tests. These comparisons were specified a priori based on our experimental design and test our core hypothesis that Martian conditions produce progressive divergence from Earth controls. Because these are planned comparisons of our primary research question rather than exploratory *post hoc* tests, we did not apply corrections for multiple comparisons. To ensure robustness of our conclusions, we verified that our strongest behavioral effects (swimming and chemotaxis) remain statistically significant when applying a conservative Holm-Bonferroni sequential correction across the three tested generations, with corrected p-values all remaining below α = 0.05. We focused our analysis on Earth versus Mars comparisons at each generation, as these directly address our hypothesis about condition effects on transgenerational trajectories. Significance was assessed at p < 0.05, and effect sizes (Cohen’s d) were calculated for all comparisons. Sample sizes for each assay are reported in figure legends and Supplementary Data 1, which contains all raw measurements for morphology, crawling kinematics, swimming frequency, and chemotaxis across all generations and conditions. Statistical verification with Excel formulas for all reported comparisons is provided in Supplementary Data 2.

We quantified trait variability by calculating coefficients of variation (CV = 100 × SD / mean) for each trait, generation and condition across replicate assays (n = 5 per group). To compare relative variability between conditions we report Mars to Earth CV ratios. Because sample sizes are small and the variability analysis is primarily descriptive, we did not perform formal hypothesis tests on differences in CV and instead focused on the magnitude and consistency of effect sizes across traits. All per assay measurements for morphology, crawling kinematics, swimming frequency and chemotaxis, together with the corresponding coefficients of variation for each trait, generation and condition, are provided in Supplementary Data 1, with statistical tests detailed in Supplementary Data 2.

### Use of AI Tools

Large language models (ChatGPT, OpenAI and Claude, Anthropic) were used to improve language and for parts of diagram in Fig. 1B. All content was subsequently checked, edited and approved by the authors.

## Results

### Morphological Changes Across Generations (Figure 2)

Earth and Mars worms followed diverging trajectories across generations, consistent with a generation by condition interaction. We did not formally test the interaction term in a two-way model, but linear fits within each condition show positive slopes for Earth and nonlinear or flat trajectories for Mars. For body length (50th percentile), Earth-grown worms showed a linear increase across generations (slope = 16.0 µm/generation) while Mars-grown worms showed a non-linear trajectory (initial increases then decreases). This differential growth pattern resulted in significant Earth versus Mars differences at Generation 2 (t(8) = 2.71, p = 0.027; Earth: 962 ± 13 µm vs. Mars: 867 ± 33 µm; Cohen’s d = 1.71) and Generation 6 (t(8) = 2.81, p = 0.02; Earth: 1010 ± 14 µm vs. Mars: 877 ± 45 µm; Cohen’s d = 1.78), but not at Generation 4 (t(8) = −1.37, p = 0.21).

**Figure 2.**
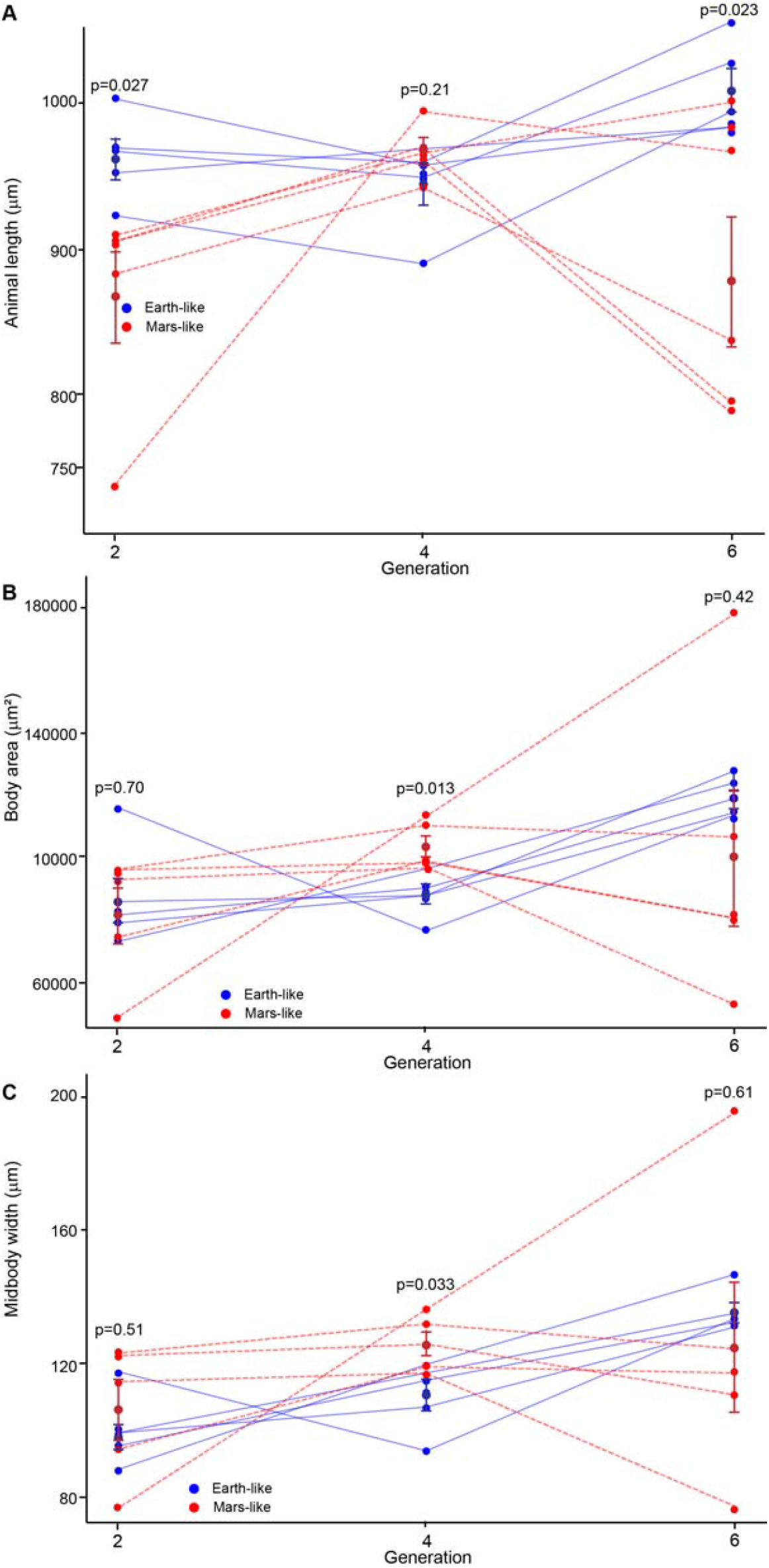
Morphological changes under Martian conditions across generations. **A)** Body length, **B)** area, and **C)** midbody width at the 50th percentile for Earth (blue circles) and Mars (red squares) conditions at Generations 2, 4, and 6. Each data point represents the mean of 5 biological replicates (15 worms per replicate plate), with error bars showing SEM. Earth and Mars worms followed diverging trajectories across generations for body length, with Earth worms showing progressive linear growth (slope = 16.0 µm per generation) and Mars worms showing a nonlinear trajectory. Earth versus Mars differences were significant at Generation 2 (t(8) = 2.71, p = 0.027) and Generation 6 (t(8) = 2.81, p = 0.023), but not Generation 4 (t(8) = −1.37, p = 0.209). Body area and midbody width showed anomalous patterns at Generation 4, where Mars-grown worms were paradoxically larger than Earth worms (Area: t(8) = −3.16, p = 0.013; Width: t(8) = −2.58, p = 0.033). Mars worms showed markedly increased phenotypic variability at Generation 6, with standard errors three to eight times greater than Earth controls, indicating developmental instability under chronic Martian conditions. These patterns demonstrate cumulative, transgenerational effects of simulated Martian gravity and magnetism on animal morphology, with evidence of non-monotonic phenotypic trajectories in Mars worms.

Body area and midbody width showed similar patterns of Generation × Condition interaction, though with different timing: Mars-grown worms paradoxically showed transiently larger body area and midbody width at Generation 4 (Area: t(8) = −3.16, p = 0.01; Earth: 88,369 ± 3,212 µm^2^ vs. Mars: 103,288 ± 3,468 µm^2^; Cohen’s d = −2.00; Width: t(8) = −2.58, p = 0.03; Earth: 110.6 ± 4.7 µm vs. Mars: 126.0 ± 3.6 µm; Cohen’s d = −1.63), suggesting a possible compensatory growth response at intermediate generations. By Generation 6, Earth-grown worms showed larger mean body area (118,648 ± 2,966 µm^2^ vs. 99,955 ± 21,526 µm^2^), though increased variability in Mars worms, reflected in standard errors that were several fold larger than in Earth controls, prevented this difference from reaching statistical significance (t(8) = 0.86, p = 0.42).

Notably, Mars worms exhibited markedly increased phenotypic variability at Generation 6 across all three traits. The coefficient of variation (CV) for Mars worms was 3.1 times higher than Earth controls for body length (5.13 % versus 1.42 %), 7.3 times higher for body area (21.53 % versus 2.50 %), and 7.0 times higher for midbody width (15.62 % versus 2.06 %). These changes indicate not only shifts in mean values but are consistent with reduced developmental precision, in which genetically identical Mars lineages produce a wider range of phenotypes despite uniform rearing conditions (Supplementary Figure S1). When compared with locomotor and sensory traits, these morphological CV increases at Generation 6 are substantially larger than the modest variability changes observed for Tierpsy kinematic metrics and the near parity in swimming and chemotaxis, as summarized in Figure 5 and Supplementary Figures 1 and 2, and the underlying values are available in Supplementary Data 1. CV profiles across the 10th, 50th and 90th percentiles further showed that median phenotypes were most affected, with 3.6 to 8.6 fold CV increases compared to 2.1 to 5.2 fold changes at the distribution extremes. This pattern suggests that typical body plans, which are normally most tightly buffered, become especially unstable under prolonged Mars like conditions.

### Swimming Frequency (Figure 3)

The crawling behavior analyzed in our morphometric assays above is not physiologically challenging, even for compromised animals. For example, dystrophic worms have been shown to display increased crawling velocity compared to healthy controls, yet they often reveal their impairments when tasked with more challenging behavioral paradigms such as burrowing or swimming (Beron et al., 2015). Consistent with this, while Tierpsy analysis revealed trends in crawling deficits for Martian conditions, these did not reach statistical significance. Tierpsy kinematic metrics that did not differ in mean between Earth and Mars conditions showed only modest variability increases (CV ratios around 1.5, Figure 5), which is intermediate between the large variability increases observed for morphology and the near parity seen for swimming and chemotaxis. To investigate whether the effect of Martian physics on locomotion became apparent under increased motor demands, we filmed and analyzed the swimming behavior of worms across different conditions. We measured swimming frequency by manually tracking the number of uninterrupted swimming cycles per second in liquid NGM.

**Figure 3.**
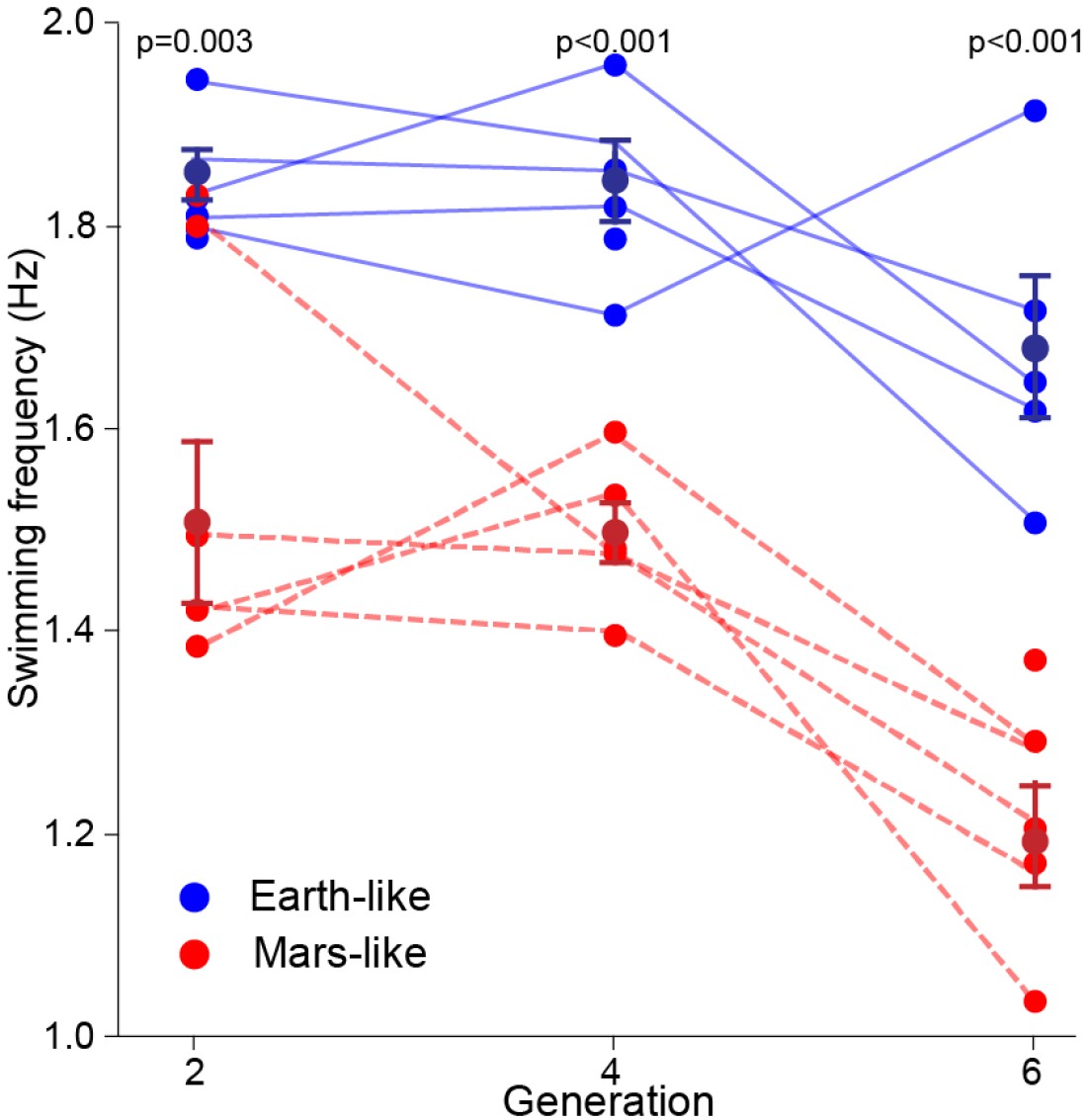
Swimming frequency under Earth vs. Mars conditions. **A)** Individual worm swimming rates (Hz) at generations 2, 4, and 6 showing distribution of values for Earth (blue) and Mars (red) conditions. **B)** Mean swimming frequency ± SEM across generations. Purple data points represent generation 0 control animals (worms raised in the experimental setup with clinostat and magnetic coils not powered). Each data point represents the mean of 5 biological lineages, with 7 worms analyzed per lineage (35 total worms per condition per generation). Planned comparisons (two-tailed t-tests) between Earth and Mars conditions at each generation showed that Mars-reared worms swam significantly slower than Earth-reared worms at all generations tested (Generation 2: p < 0.05; Generation 4: p <0.05; Generation 6: p < 0.05). Earth worms maintained stable swimming frequency (1.852 ± 0.026 Hz to 1.682 ± 0.069 Hz) across all generations, while Mars worms exhibited progressive decline from 1.511 ± 0.078 Hz (Generation 2) to 1.196 ± 0.047 Hz (Generation 6).

Swimming frequency was measured at Generations 2, 4, and 6 to assess locomotor function under energetically demanding conditions. Mars-reared worms showed significantly reduced swimming frequency compared to Earth controls at all generations tested. At Generation 2, Mars worms swam at 1.511 ± 0.08 Hz compared to Earth worms at 1.85 ± 0.03 Hz (t(8) = 4.17, p = 0.003, Cohen’s d = 2.64). This impairment persisted at Generation 4 (Mars: 1.500 ± 0.03 Hz vs Earth: 1.849 ± 0.04 Hz; t(8) = 6.66, p < 0.001, Cohen’s d = 4.21) and Generation 6 (Mars: 1.20 ± 0.05 Hz vs Earth: 1.68 ± 0.07 Hz; t(8) = 5.86, p < 0.001, Cohen’s d = 3.70). Notably, while Earth worms maintained relatively stable swimming frequency across generations, Mars worms exhibited progressive decline, with Generation 6 showing a 29% reduction in swimming frequency relative to Earth controls, indicating sustained impairment of high-energy locomotor circuits under simulated Martian conditions.

### Neurological Function (Figure 4)

Chemotaxis toward diacetyl was assessed at Generations 2, 4, and 6 to evaluate sensory-neural function. Mars-reared worms showed comparable chemotaxis indices to Earth controls at Generation 2 (Earth: 0.96 ± 0.02 vs Mars: 0.91 ± 0.03; t(8) = 1.48, p = 0.177), indicating intact sensory function at early generations. However, significant deficits emerged by Generation 4 (Earth: 0.99 ± 0.01 vs Mars: 0.89 ± 0.03; t(8) = 2.79, p = 0.02, Cohen’s d = 1.77) and became more pronounced at Generation 6 (Earth: 0.91 ± 0.03 vs Mars: 0.72 ± 0.02; t(8) = 5.82, p < 0.001, Cohen’s d = 3.68). The progressive decline in chemotactic ability under Martian conditions suggests that chronic hypogravity and hypomagnetic stress produce cumulative transgenerational effects on sensory neuron function or downstream circuit integrity.

**Figure 4.**
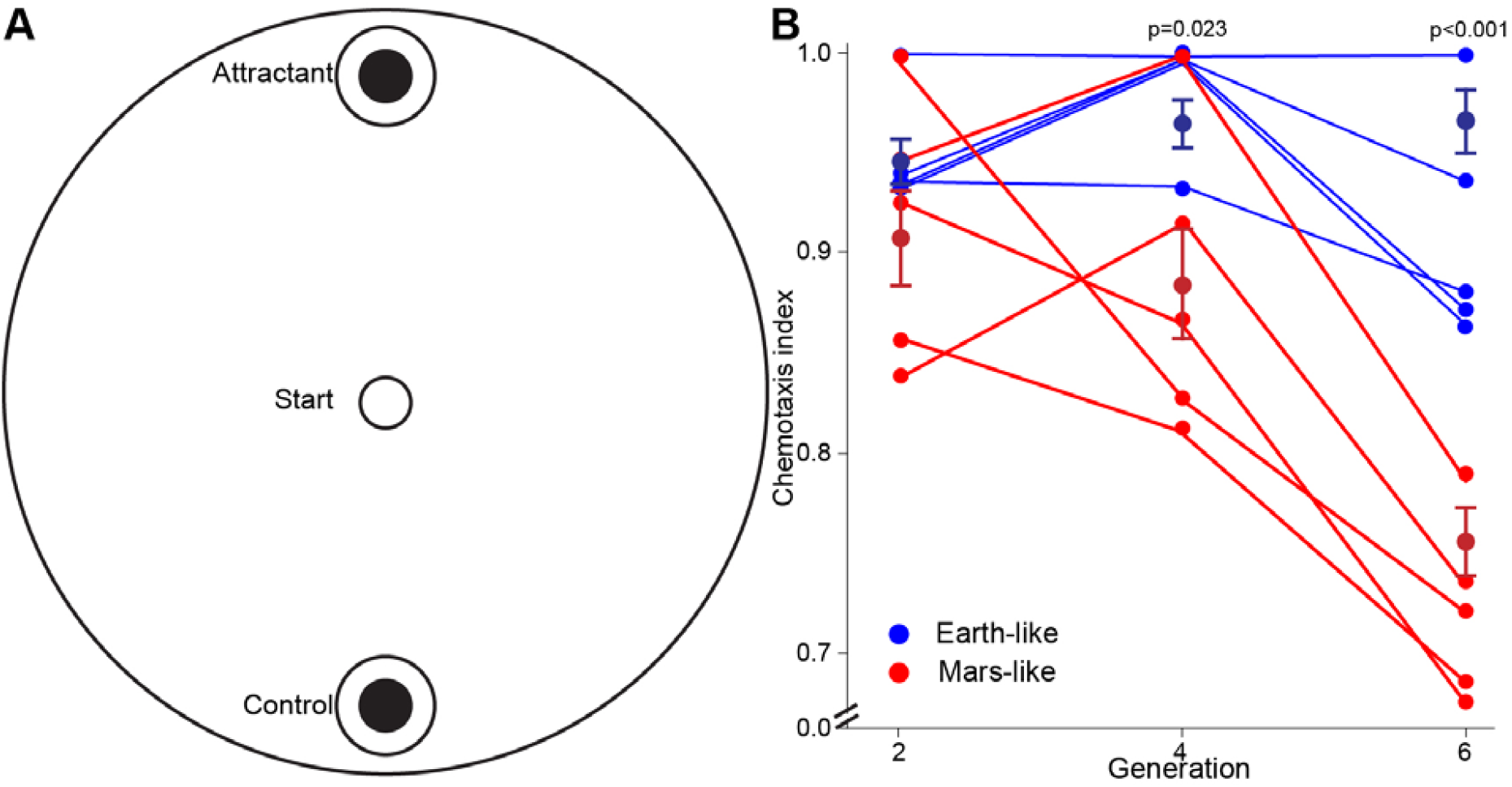
Chemotaxis index over generations. **A)** Schematic of chemotaxis assay. Animals were placed at the center of a 10cm agar plate with chemotaxis agar. A drop of diacetyl attractant (1×10^−3^M) was placed on one side and a control drop of NGM on the opposite side. Animals were allowed to freely crawl on the plate and worms immobilized at either test or control spots were tallied after 30 minutes. **B)** Chemotaxis indices for animals raised under control conditions (G0, purple) or under Earth (blue) vs. Mars (red) conditions for 2, 4, and 6 generations. N = 5 biological lineages; error bars = SEM. Planned comparisons (two-tailed t-tests) between Earth and Mars conditions at each generation showed that Mars worms had chemotaxis indices comparable to Earth controls at Generation 2 (Earth: 0.963 ± 0.015 vs Mars: 0.914 ± 0.029; t(8) = 1.48, p = 0.177) but exhibited significant deficits by Generation 4 (Earth: 0.987 ± 0.013 vs Mars: 0.885 ± 0.034; t(8) = 2.79, p = 0.023) and Generation 6 (Earth: 0.911 ± 0.026 vs Mars: 0.722 ± 0.020; t(8) = 5.82, p < 0.001), demonstrating progressive sensory decline under Martian conditions.

**Figure 5.**
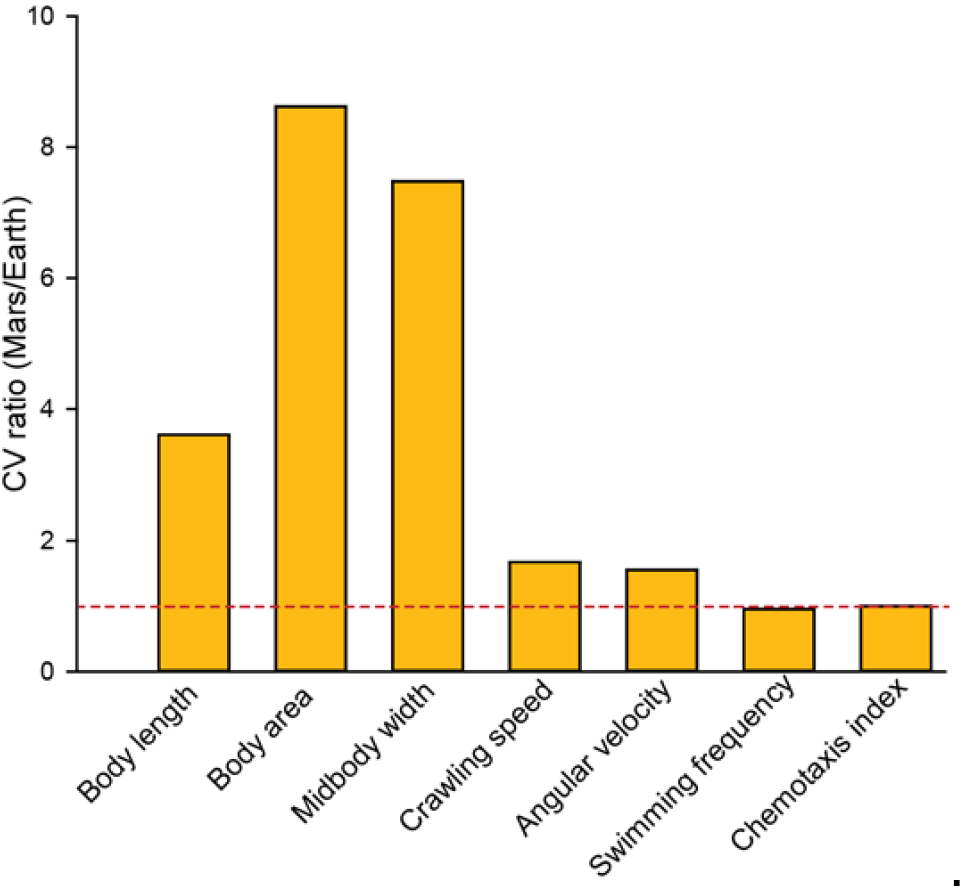
Trait specific variability at generation six under Mars like conditions. Bars show the ratio of coefficients of variation (CV Mars divided by CV Earth) at generation six for morphological traits (body length, body area, midbody width), Tierpsy kinematic measures (crawling speed, angular velocity), and behavioral performance traits (swimming frequency, chemotaxis index). CVs were calculated as one hundred times standard deviation divided by mean across replicate assays (n = 5 plates per condition). The dashed line marks equal variability for Mars and Earth. Morphological traits show three to eight fold higher variability under Mars conditions, Tierpsy kinematic measures show modest increases around one point six fold, and swimming and chemotaxis show CV ratios near one despite clear mean performance deficits.

## Discussion

Organisms raised under simulated Martian gravity and magnetism remained viable across multiple generations but developed a range of physiological impairments that emerged gradually and differed in severity across functional domains. While early generations exhibited relatively subtle changes, by the sixth generation, clear divergence emerged between Earth and Mars cohorts across behavioral, morphological, and sensory measures. These findings reveal that the effects of Martian physics are cumulative, producing transgenerational stress phenotypes that may not be detectable in shorter-term exposures.

A critical question is whether the combined Martian conditions we tested produce effects distinct from either factor alone. Our study did not include single parameter controls (gravity alone or magnetism alone), so we cannot quantitatively apportion the observed changes to gravity versus magnetism. Our experimental design was therefore intentionally not factorial. Here our goal was different: to treat Martian gravity and magnetism as coupled aspects of a single planetary environment and to ask whether that environment, taken as a whole, can drive multigenerational phenotypic change in an Earth-evolved lineage. This planet centric perspective is relevant not only for Mars but for any future destination where global physical parameters, such as surface gravity and the ambient magnetic field, will inevitably differ from those on Earth. Studies of *C. elegans* in microgravity alone report decreased body length, muscle atrophy, reduced expression of muscle and cytoskeletal proteins, and locomotor deficits that parallel our observations (Selch et al., 2008; Çelen et al., 2023; Higashibata et al., 2016). Notably, space-flown *C. elegans* exhibit significant epigenetic modifications mediated by histone deacetylase HDA 4, with body length decreases becoming more pronounced in *hda 4* mutants and morphological changes accumulating across generations (Higashibata et al., 2021). By contrast, studies examining hypomagnetic field (HMF) exposure alone in *C. elegans* have reported mixed and sometimes paradoxical results. While hypomagnetic fields (HMF) can induce oxidative stress and disrupt redox homeostasis in various animal models (Tian et al., 2022, 2024), several studies and reviews indicate that HMF can also elicit context⍰dependent adaptive responses, including enhanced stress resistance, consistent with hormesis in some systems (Binhi et al., 2017). This contrasts with our finding of progressive functional decline under combined Mars like conditions and raises the possibility that the simultaneous presence of reduced gravity and near absent magnetism could produce non additive effects, although this remains speculative in the absence of single factor controls. The morphological stunting, locomotor decline and sensory deficits we observed by Generation 6 are consistent with the idea that combined stressors may overwhelm compensatory mechanisms that can partially protect against individual factors. Future studies incorporating factorial designs with isolated gravity and magnetic field manipulations will be essential to determine whether Mars like conditions produce synergistic, antagonistic or additive effects on biological function.

### Morphological divergence and compensatory responses

Our analysis (Figure 2) revealed complex, non-monotonic morphological trajectories under Martian conditions. For body length, we observed significant Earth versus Mars divergence at Generation 2 (t(8) = 2.71, p = 0.027) and most prominently at Generation 6 (t(8) = 2.81, p = 0.023), but not at Generation 4. Most strikingly, Mars worms at Generation 4 showed paradoxically larger body area (17% increase, t(8) = −3.16, p = 0.013, Cohen’s d = −2.00) and midbody width (14% increase, t(8) = −2.58, p = 0.033, Cohen’s d = −1.63) compared to Earth controls. Rather than representing an anomaly, this pattern is consistent with hormesis, a well-documented biological phenomenon where moderate chronic stress triggers beneficial adaptive responses. In *C. elegans*, environmental stressors can induce transgenerationally inheritable survival advantages, with exposed parents showing initial impairment but their descendants exhibiting enhanced stress resistance that persists for multiple generations through epigenetic mechanisms involving DAF-16/FOXO and HSF-1 signaling pathways (Kishimoto et al., 2017). Meta analysis of 198 effect sizes supports the interpretation that hormesis in C. elegans enhances performance metrics by 16 to 25% (Wang et al., 2020), closely matching our observed morphological increases at Generation 4. Notably, Jobson et al. (2015) demonstrated that while starved *C. elegans* larvae exhibit reduced growth and fertility, their immediate descendants show increased resistance to subsequent starvation and heat stress, directly paralleling our observation that Mars worms transition from initial morphological deficits at Generation 2 to compensatory overgrowth at Generation 4. This pattern is consistent with the idea that Martian conditions may elicit a hormetic response that builds across generations, reaching maximum compensatory capacity by the fourth generation before ultimately declining. Because we did not directly measure survival or stress resistance, we cannot determine whether this represents true hormesis in the strict sense or a transient plastic overgrowth response.

The non-monotonic morphological trajectory we observe, with an initial deficit at Generation 2, a transient increase at Generation 4, and subsequent decline by Generation 6, differs from simple models of progressive deterioration under stress. Similar peak-and-decline temporal profiles have been described for epigenetically mediated traits in *C. elegans*. For example, Greer and colleagues reported that longevity benefits from altered H3K4 methylation are largest in F3 and F4 descendants before fading in later generations (Greer et al. 2011), and Klosin and colleagues showed that chromatin changes induced by temperature can persist for many generations before returning toward the ancestral state (Klosin et al. 2017). These studies illustrate that multigenerational environmental responses can include intermediate generations with enhanced or compensated phenotypes. Our data are consistent with this type of transient compensation, although direct molecular measurements will be required to determine whether comparable epigenetic mechanisms underlie the morphological changes we observe.

### Molecular mechanisms underlying transgenerational responses

Several molecular pathways likely contribute to the transgenerational compensatory response we observe. The molecular basis for such transgenerational compensation likely involves small RNA communication systems, as recent work demonstrates that miRNA-processing enzymes (DRSH-1) and worm-specific Argonaute proteins are essential for both the acquisition and inheritance of hormesis effects through neuron-to-intestine-to-germline signaling pathways (Okabe et al., 2021). The insulin/IGF signaling (IIS) pathway, which regulates both stress resistance and developmental programs in *C. elegans*, is known to be altered by microgravity and plays a central role in hormetic responses, with *daf-16/FOXO* and *daf-18* mutations blocking hormetically-induced adaptations (Selch et al., 2008; Cypser and Johnson, 2006). Transgenerational inheritance of stress-adaptive phenotypes requires germline-expressed small RNAs and the nuclear RNAi machinery, including the germline Argonaute HRDE-1 complex, which transmits epigenetic information across generations through chromatin modifications including H3K4 and H3K36 methylation (Rechavi et al., 2014). The progressive accumulation of compensatory signals from Generation 2 to Generation 4 suggests that epigenetic marks build across successive germline transmissions, analogous to the multi-generation accumulation of chromatin changes observed in temperature stress studies (Klosin et al., 2017). However, the metabolic cost of maintaining compensatory responses, combined with progressive decay of epigenetic signals through active “forgetting” mechanisms, likely explains why compensation peaks at Generation 4 rather than continuing indefinitely. The ultimate failure of buffering capacity by Generation 6, manifested as dramatically increased phenotypic variability, is consistent with depletion of epigenetic regulatory systems such as histone deacetylase HDA-4, which previous spaceflight studies have identified as critical for buffering developmental stress (Higashibata et al., 2021).

### Developmental canalization and the loss of phenotypic buffering

The most revealing aspect of the morphological data is the pattern of phenotypic variability across generations. By Generation 6, Mars-reared lineages showed 3.1 fold higher CV for body length, 7.3 fold higher CV for body area, and 7.0 fold higher CV for midbody width than Earth controls. These large variance differences, in the absence of any change in genotype or rearing protocol, are consistent with reduced developmental canalization, meaning a decreased ability of developmental programs to produce similar phenotypes under perturbation (Debat and David, 2001; Flatt, 2005). While we did not conduct formal statistical tests on variance differences given our sample sizes, the three-to eight-fold CV increases represent large descriptive effect sizes that are consistent with substantially reduced phenotypic buffering.

In our measurements, morphology, which is set during development, showed increasing variance over generations, whereas adult performance traits such as swimming and chemotaxis converged toward uniformly impaired but relatively consistent values. If similar distinctions hold for other traits, then developmental processes may be more susceptible to loss of predictability under Mars like conditions than some adult behaviors.

CV profiles across the morphological distribution revealed that median phenotypes were most affected, with 3.6 to 8.6 fold increases in variability at the 50th percentile compared with smaller changes at the 10th and 90th percentiles. One possible explanation is that individuals at the extremes already operate closer to biomechanical or physiological limits, which may restrict further divergence, whereas typical body plans have more scope to vary once buffering mechanisms are weakened. This interpretation aligns with theoretical work on canalization and developmental stability, in which stabilizing selection is thought to act most strongly on common phenotypes (Flatt, 2005).

Several mechanisms could underlie the reduced canalization we observe. Spaceflight studies in *C. elegans* implicate epigenetic regulators such as the histone deacetylase HDA 4 in buffering microgravity effects, and mutants lacking this regulator show exaggerated morphological defects under spaceflight like conditions (Higashibata et al., 2021). More generally, work on stress induced plasticity has shown that when canonical buffers like HSF 1 are compromised, unfolded protein response and innate immunity pathways can partially compensate, suggesting a broader network of developmental stabilizers (Kovács et al., 2024).

### Locomotor vulnerability under energetic challenge

While our high-throughput Tierpsy crawling analysis revealed trends toward reduced crawling performance in Mars animals, these did not reach statistical significance. This may reflect the relatively low energy demands of crawling, which even physiologically impaired animals can often maintain. In contrast, our manual swimming assay revealed consistent and significant locomotor impairments in Mars worms at every generation tested (Figure 3). Swimming requires sustained, high-frequency neuromuscular activation and coordination, making it more sensitive to subtle neuromotor deficits. The progressive decline in swimming frequency among Mars worms (from 1.511 ± 0.078 Hz at Generation 2 to 1.20 ± 0.05 Hz at Generation 6), despite stable performance in Earth controls (1.85 ± 0.03 Hz to 1.68 ± 0.07 Hz), suggests that the Martian environment impairs energy-intensive motor circuits in a cumulative, dose-dependent manner.

This pattern mirrors results from studies of dystrophic or mitochondrial mutants, where stress only becomes apparent under biomechanical challenge (Çelen et al., 2023; Szewczyk et al., 2005). It also parallels the spaceflight-induced downregulation of muscle-associated genes observed in *C. elegans* and mammals (Selch et al., 2008; Morey-Holton, 2003). Microgravity alone is sufficient to cause decreased myosin heavy chain expression, reduced muscle contractility, and significant motility abnormalities in *C. elegans* (Higashibata et al., 2016), effects that persist across multiple generations and mirror the muscle atrophy seen in astronauts (Moosavi et al., 2021). The swimming deficits we observed are consistent with this microgravity literature, but the consistent impairment across all generations tested (rather than adaptation) and the magnitude of decline by Generation 6 suggest that combined stressors may prevent the partial compensation sometimes observed in microgravity-only exposures. Whether hypomagnetic conditions independently contribute to neuromuscular dysfunction or simply fail to provide protective effects remains unclear and warrants targeted investigation.

### Sensory decline across generations

In addition to motor impairments, we observed a transgenerational decline in chemotaxis performance under Martian conditions (Figure 4). Mars worms began with chemotaxis indices comparable to Earth controls at Generation 2 but showed significant deficits by the fourth generation (t(8) = 2.79, p = 0.023), which deepened further by the sixth generation (t(8) = 5.82, p < 0.001). Chemotaxis toward diacetyl depends on the function of AWA and AWC sensory neurons, as well as downstream interneurons involved in directional navigation (Bargmann, 2006). These neurons are known to rely on redox-sensitive signaling pathways that may be disrupted by chronic hypomagnetic conditions.

The observed decline in sensory performance may reflect multiple mechanisms. Hypomagnetic fields have been shown to interfere with ROS homeostasis, neural excitability, and behavioral responses across species (Tian et al., 2022, 2024; Zhang and Tian, 2020). The progressive nature of the chemotaxis deficit is particularly intriguing given that short⍰term HMF exposure has been reported, in some contexts, to increase stress resistance rather than impair it, consistent with hormetic responses to hypomagnetic conditions (Bihni et al., 2017). This suggests that while acute HMF exposure may trigger beneficial hormetic responses, chronic multigenerational exposure, particularly in combination with reduced gravity, leads to cumulative dysfunction. Oxidative stress or impaired radical-pair mechanisms, as seen in models of cryptochrome-mediated magnetoreception (Ritz et al., 2000), could contribute to this progressive sensory decline. Alternatively, the combination of reduced proprioceptive input from hypogravity and altered redox signaling from HMF may synergistically disrupt the integration of sensory information required for effective chemotaxis.

### Mechanosensory and proprioceptive constraints

Postural simplification in Mars-reared worms may reflect biomechanical unloading, reduced proprioceptive input, or altered neuronal tone. In *C. elegans*, postural control involves mechanosensory feedback through gravity-sensitive channels such as PEZO-1 and associated proprioceptors (Ackley et al., 2025). The flattening observed in curvature metrics over time may therefore represent either an adaptive energy-conserving response or a breakdown in sensory-motor integration under long-term gravitational and magnetic deprivation.

### Differential vulnerability across functional domains

The distinct temporal patterns of impairment across functional domains reveal differential vulnerability to Martian conditions. Swimming, which requires sustained high-energy neuromuscular output, showed immediate and severe impairment at Generation 2 (Cohen’s d = 2.64) that persisted across all generations tested, with the largest effect sizes observed in this study (d = 4.21 at Generation 4). In contrast, chemotaxis remained comparable to Earth controls at Generation 2 but showed progressive decline by Generations 4 (d = 1.77) and 6 (d = 3.68), suggesting that sensory systems may possess greater initial resilience or buffering capacity. Morphological traits exhibited yet another pattern, with non-monotonic trajectories showing initial deficit, transient compensatory growth at Generation 4, and ultimate decline with high variability by Generation 6. This pattern indicates that energy-intensive motor functions represent the most vulnerable biological systems under chronic Martian stress, with implications for astronaut physical work capacity and exercise tolerance during long-duration Mars missions. The eventual decline of motor, sensory, and developmental systems by Generation 6 suggests that while timing varies by functional domain, no biological system is spared from the cumulative effects of sustained Martian conditions.

### Transgenerational Mechanisms: Epigenetic vs. Genomic Inheritance

The progressive nature of the phenotypes we observed, with impairments becoming most pronounced by Generation 6, raises important questions about the mechanisms underlying transgenerational effects. Several non-exclusive possibilities exist: epigenetic inheritance, accumulated cellular damage, genetic mutations, or adaptive selection.

Epigenetic mechanisms are well-documented in *C. elegans* and represent a plausible pathway for transgenerational responses to environmental stress. *C. elegans* can transmit environmentally induced changes through small RNAs, histone modifications, and chromatin remodeling, with effects persisting for multiple generations (Baugh and Day, 2020). Temperature-induced chromatin changes in *C. elegans* can persist for at least 14 generations through both maternal and paternal inheritance (Klosin et al., 2017), and starvation stress induces transgenerational effects on growth, reproduction, and stress resistance mediated by small RNAs and epigenetic pathways (Rechavi et al., 2014; Jobson et al., 2015). Similarly, learned pathogen avoidance behavior can be transmitted through multiple generations via Piwi/PRG-1 argonaute-dependent small RNA pathways (Moore et al., 2019; Akinosho et al., 2025), demonstrating that behaviorally-relevant environmental information can be inherited through germline epigenetic mechanisms. Notably, space-flown *C. elegans* show epigenetic modifications via histone deacetylase HDA-4, with mutants lacking this epigenetic regulator showing exacerbated morphological defects under microgravity (Higashibata et al., 2021). This suggests that epigenetic buffering normally mitigates some spaceflight effects, and that its dysregulation or saturation over multiple generations could explain the progressive phenotypes we observed.

Alternatively, or additionally, accumulated cellular damage and impaired repair mechanisms could compound across generations. Oxidative stress from hypomagnetic conditions (Tian et al., 2022, 2024) combined with reduced mechanical stimulation from hypogravity may progressively damage proteins, lipids, and DNA. If repair pathways are compromised under Martian conditions, as suggested by altered mitochondrial function in space-flown organisms (Higashibata et al., 2016), damage could accumulate both within and across generations. The progressive swimming decline and sensory deficits we observed are consistent with cumulative cellular dysfunction.

Genetic mutation and selection represent a third possibility, though less likely over only six generations. While space radiation is known to be mutagenic, our ground-based study lacked radiation exposure. It is conceivable that altered redox environments or DNA repair deficiencies in Martian-like conditions could elevate mutation rates. However, true genetic adaptation typically requires many more generations and would be expected to produce improved rather than declining fitness under the selecting conditions.

Most likely, the transgenerational effects we observed reflect a combination of these mechanisms, with epigenetic changes and accumulated damage playing primary roles. Future work incorporating transcriptomic, proteomic, and chromatin accessibility assays across generations, as well as examination of small RNA inheritance and histone modification patterns, will be essential to dissect the relative contributions of these pathways. Understanding these mechanisms is critical not only for predicting long-term biological responses to Martian habitation but also for developing targeted interventions to mitigate transgenerational decline.

### Implications for Martian colonization

Together, these findings show that Mars like gravity and magnetism are sufficient to produce cumulative heritable changes in a terrestrial animal model. The selective sensitivity of different traits in our study, including morphology, locomotion and chemotaxis, suggests that some biological systems may be more vulnerable than others to altered planetary physics. Many of the phenotypes became clearly different from Earth controls only after several generations, which emphasizes the importance of multigenerational experiments when considering long term habitation scenarios.

A central implication of our data concerns developmental predictability. By Generation 6, Mars lineages showed three to eight fold increases in morphological variability, indicating disruption of the homeostatic processes that usually keep development within a narrow range. If comparable effects occurred in crops, livestock or humans, they could in principle complicate reliable agriculture, medical care and population management even in situations where mean trait values remained within acceptable limits. We therefore suggest that maintaining developmental canalization may be as important as preserving average performance when designing biological countermeasures for sustained life on Mars.

### Caveats and limitations

While our study offers insight into the long-term impact of Martian-like gravity and magnetism on a model organism, several limitations should be acknowledged. First, although our clinostat and Merritt cage setup effectively simulates reduced gravity and attenuated magnetic fields, these are ground-based approximations. The clinostat cannot reproduce true microgravity and is most effective at long time scales. It may thus not fully mimic low-gravity biomechanics relevant to fast locomotion or rapid intracellular transport. Similarly, our use of homogeneous magnetic field cancellation does not reflect the patchy crustal magnetism found on Mars, which may variably impact orientation or neural function.

Second, we used only a single laboratory strain (N2) of *C. elegans*. Natural isolates or genetically diverse lines may exhibit different sensitivity or adaptive trajectories under Martian analog conditions. The N2 strain has been laboratory-adapted for decades and may not represent the full range of phenotypic plasticity present in wild *C. elegans* populations.

Third, as noted above, our design did not include single-parameter controls that would allow us to definitively determine whether the observed effects result from reduced gravity alone, hypomagnetic conditions alone, or their interaction. While we have contextualized our findings against published literature examining these factors independently, factorial experimental designs with isolated manipulations of gravity and magnetic field will be necessary to establish whether Mars like conditions produce synergistic, antagonistic, or simply additive effects. Such studies would clarify whether the progressive decline we observed represents a unique signature of combined stressors or simply reflects the sum of individual factor effects.

Finally, our study focused on phenotypic outcomes without probing the mechanistic basis of these changes. While we have proposed several plausible mechanisms (epigenetic inheritance, accumulated damage, impaired repair), direct evidence is lacking. The use of high-throughput transcriptomic, proteomic, and chromatin accessibility assays could help differentiate between gene regulatory changes, epigenetic inheritance, and cumulative stress. Notably, we did not analyze embryogenesis, a developmental stage likely to be highly sensitive to altered physical fields. Future work should examine embryonic development in detail, including lineage progression, axis formation, and cell cycle timing, as potential loci of environmentally induced, heritable change.

### Future directions

Several key experimental directions emerge from this work. First, to establish the relative contributions of gravity and magnetism to the phenotypes we observed, factorial designs with four conditions (Earth gravity + Earth magnetism, Earth gravity + Mars magnetism, Mars gravity + Earth magnetism, and Mars gravity + Mars magnetism) should be conducted. Such experiments would definitively determine whether combined Mars conditions produce non-additive effects and identify which biological systems are most sensitive to each factor.

Second, to pinpoint molecular underpinnings, transcriptomic and proteomic profiling across generations should be conducted to reveal pathways such as insulin/IGF-1 (Çelen et al., 2023), TGF-β (Selch et al., 2008), and iron homeostasis (Rouault, 2013) that are altered by hypogravity and hypomagnetic fields. Small RNA sequencing and chromatin immunoprecipitation could identify epigenetic marks that mediate transgenerational inheritance. Comparison of wild-type animals with mutants lacking key epigenetic regulators (such as *hda-4, hrde-1*, or components of the piRNA pathway) would clarify whether epigenetic mechanisms are necessary for the progressive phenotypes we observed or whether other mechanisms predominate.

Third, testing interventions could inform countermeasures for long-term space habitation. Candidate interventions include cryptochrome agonists to restore magnetic field signaling, ROS scavengers to mitigate oxidative stress from hypomagnetic conditions, mechanical stimulation to compensate for reduced proprioceptive input, and exercise paradigms to maintain neuromuscular function. If specific molecular pathways are identified as primary drivers of decline, targeted pharmacological or genetic interventions could be designed and tested.

Fourth, in vivo imaging of embryogenesis under Martian analog conditions could clarify when and where morphological or behavioral changes originate. Time-lapse microscopy of embryonic cell divisions, lineage progression, and axis formation would reveal whether Martian conditions disrupt early patterning events or whether effects accumulate primarily during post-embryonic development. Given that embryogenesis represents a critical window for environmental programming, such studies could identify the most vulnerable developmental stages.

Fifth, extending these studies to natural isolates of *C. elegans* and ultimately to mammalian models will be critical to bridge the gap from nematodes to humans in future planetary missions. Natural isolates may reveal genetic variation in susceptibility to Martian conditions, potentially identifying protective alleles that could inform selective breeding or genetic engineering strategies. Mammalian studies, while more resource-intensive, are essential for validating findings in systems with greater physiological complexity and direct relevance to human health.

Finally, comparative studies examining organisms that have evolved in extreme environments on Earth (such as cave-dwelling species adapted to near-zero magnetic fields or aquatic organisms experiencing altered effective gravity) could provide insight into natural mechanisms of adaptation to conditions analogous to those on Mars. Such comparative approaches could reveal convergent solutions to similar environmental challenges and inspire biomimetic strategies for enhancing resilience to Martian conditions.

---

**Supplementary Figure S1.**
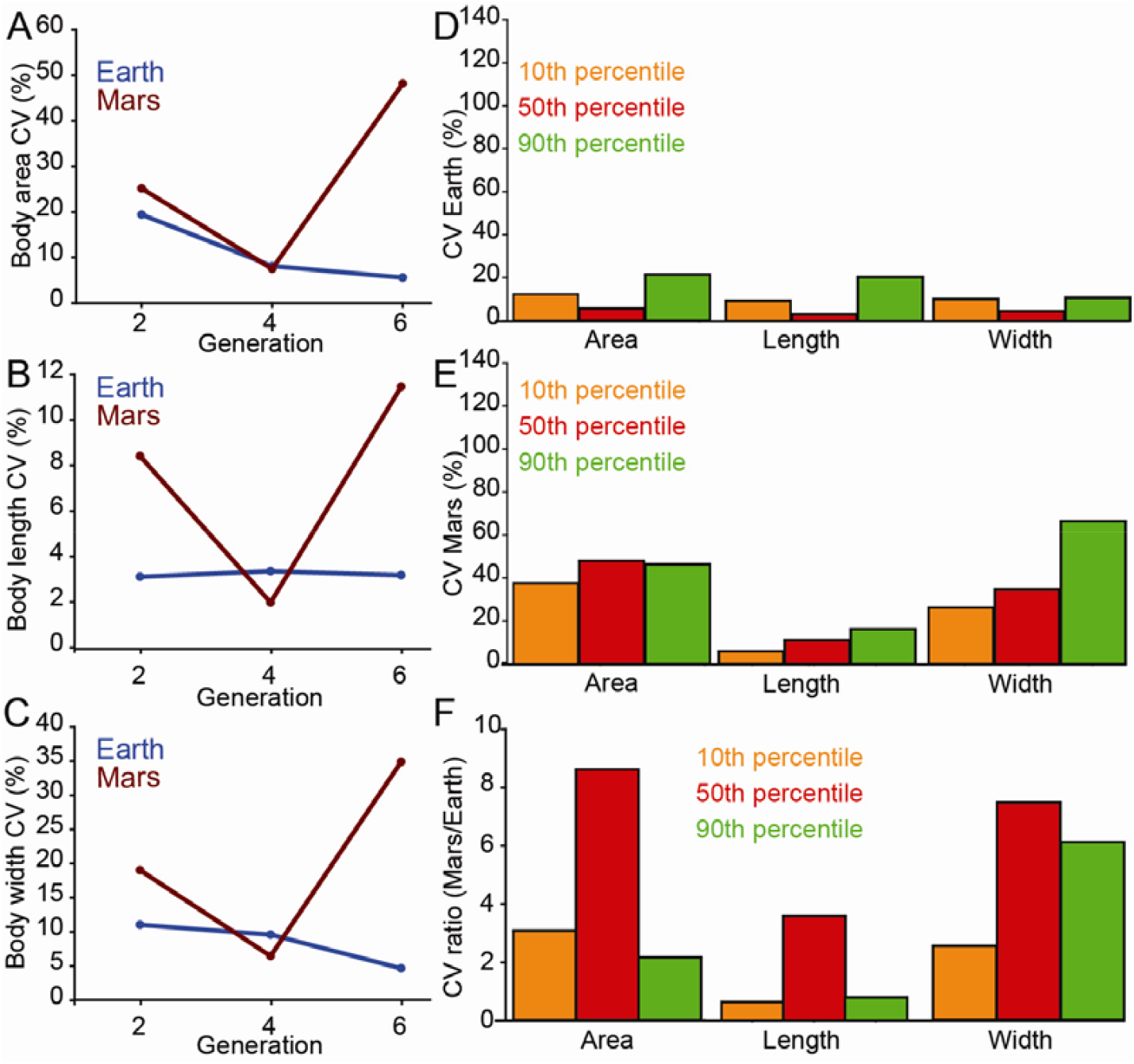
Developmental traits show increased phenotypic variability under Mars like conditions. **(A–C)** Coefficients of variation (CV, 100 × SD ÷ mean) for body area **(A)**, body length **(B)** and midbody width **(C)** in Earth (blue) and Mars (red) lineages across generations 2, 4 and 6 (n = 5 plates per condition and generation). Mars worms show a marked increase in CV by Generation 6 for all three traits, while Earth controls remain relatively stable. **(D**,**E)** CV values at Generation 6 for the 10th (orange), 50th (red) and 90th (green) percentiles of each trait on Earth **(D)** and Mars **(E). (F)** Mars/Earth CV ratios at Generation 6 for the same percentiles. Median phenotypes show the largest variability increases (up to ∼9-fold for body area and ∼7-fold for midbody width), whereas individuals at the extremes exhibit smaller or no increases. These data indicate that prolonged exposure to Mars like conditions reduces developmental precision and particularly destabilizes typical body plans.

**Supplementary Figure S2.**
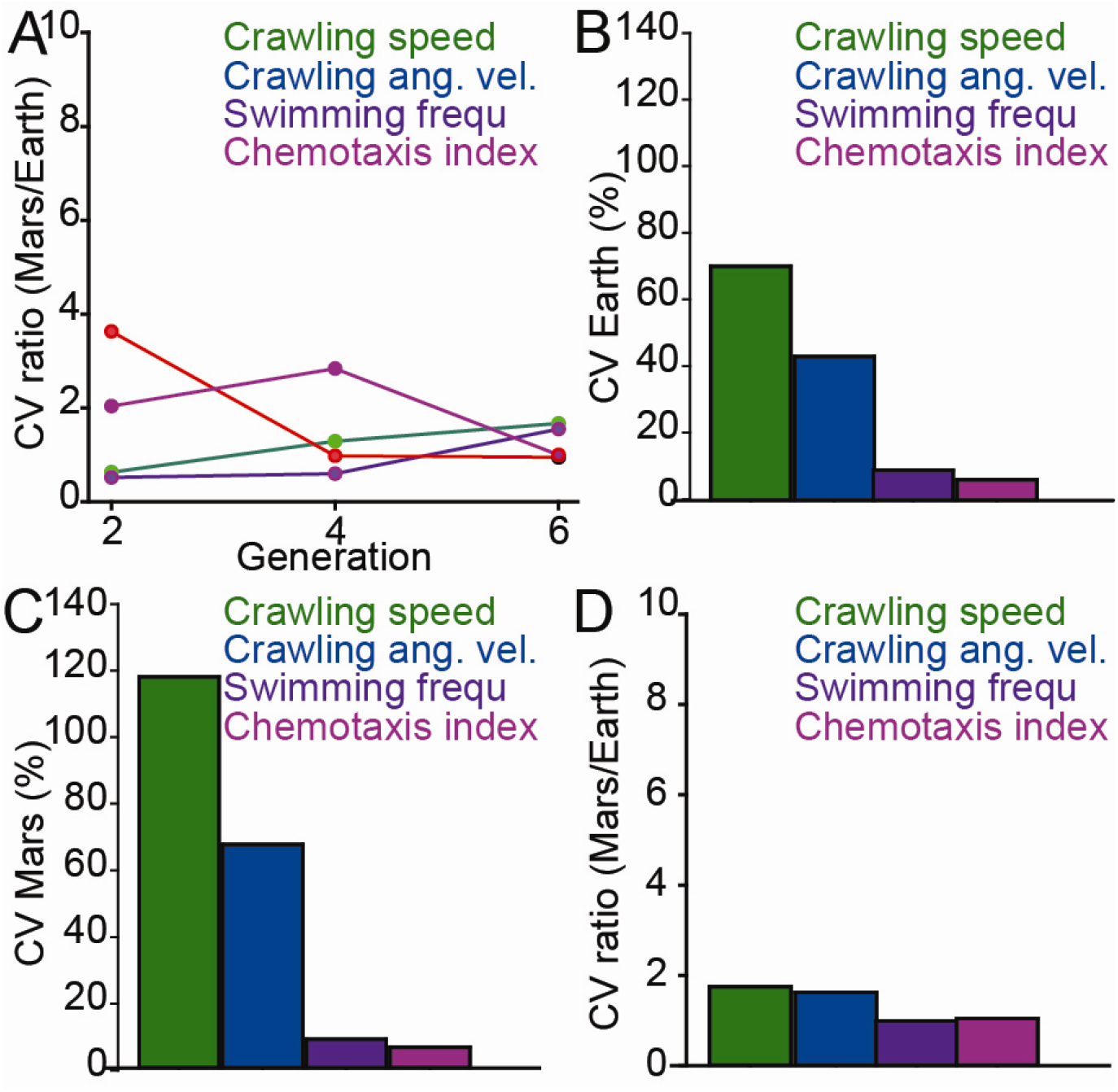
Variability in locomotor and performance traits under Mars like conditions. CV was calculated as 100 × SD ÷ mean across replicate assays (n = 5 plates per condition and generation). **A)** Mars/Earth CV ratios across generations for crawling speed, crawling angular velocity, swimming frequency and chemotaxis index. The dashed line at 1 indicates equal variability between conditions. **B)** CV values (%) at Generation 6 for the same traits on Earth. **C)** CV values (%) at Generation 6 for the same traits on Mars. **D)** Mars/Earth CV ratios at Generation 6 for crawling speed, crawling angular velocity, swimming frequency and chemotaxis index. Together with Supplementary Figure S1, these data show that locomotor kinematics exhibit modest increases in variability under Mars like conditions, whereas swimming and chemotaxis display transient early variability that returns to near parity by Generation 6.

## Data Availability

All data generated or analyzed during this study are included in this published article and its supplementary information files. Source data for Figures 1 to 5 and Supplementary Figures 1 and 2 are provided in Supplementary Data 1. Statistical analysis code is provided in Supplementary Software 1 (Python script), with independent verification workbook containing Excel formulas for all statistical tests in Supplementary Data 2. All materials enable complete reproduction of reported analyses.

## Acknowledgements

We thank members of the Vidal-Gadea laboratory for technical assistance and discussion. This work was supported by NIAMS award 2R15AR068583-02 to A.G.V-G.

## Author contributions

A.A., Z.B., X.D., W.S. and A.G.V-G. conceived the study and designed the experiments. A.A., and Z.B. performed the behavioral and morphological experiments. X.D. carried out the statistical modelling and assisted with data analysis. W.S. provided input on experimental design and interpretation of locomotor phenotypes. A.A. and A.G.V-G. analyzed the data and prepared the figures. A.A. drafted the manuscript with input from A.G.V-G. All authors reviewed and approved the final manuscript.

