## Supplementary data 3 for "Born on Mars: Multigenerational phenotypic change in *Caenorhabditis elegans* under Martian analog gravity and hypomagnetic fields": Supplementary Software 1.py

```
#!/usr/bin/env python3
"""
Statistical Analysis for Born on Mars Manuscript
================================================

This script performs all statistical analyses reported in the manuscript:
"Born on Mars: Multigenerational phenotypic change in Caenorhabditis elegans 
under Martian analog gravity and hypomagnetic fields"

Authors: Akinosho A, Benefield Z, Fritz A, Dai X, Stein W, Vidal-Gadea AG

Data source: Supplementary Data 1.xlsx
Output: Supplementary Data 2.xlsx

Requirements:
    - Python 3.8+
    - pandas
    - numpy
    - scipy
    - openpyxl
    - xlsxwriter

Usage:
    python Supplementary_Software_1.py

This will:
1. Load data from Supplementary Data 1.xlsx
2. Perform all statistical tests (t-tests, effect sizes)
3. Generate Supplementary Data 2.xlsx with formulas
4. Print summary of all results
"""

import pandas as pd
import numpy as np
from scipy import stats
from pathlib import Path
import sys

# Configuration
INPUT_FILE = "Supplementary Data 1.xlsx"
OUTPUT_FILE = "Supplementary Data 2.xlsx"
ALPHA = 0.05  # Significance level

def load_data(filepath):
    """
    Load data from Excel file.
    
    Parameters:
        filepath (str): Path to Supplementary Data 1.xlsx
        
    Returns:
        dict: Dictionary with keys 'morphology', 'swimming', 'chemotaxis'
    """
    print(f"Loading data from {filepath}...")
    
    data = {}
    
    # Load morphology data (Figure 2)
    df_morph = pd.read_excel(filepath, sheet_name='Figure 2')
    # Filter to generations 2, 4, 6 and earth/mars only
    df_morph = df_morph[df_morph['Generation'].isin([2.0, 4.0, 6.0])]
    df_morph = df_morph[df_morph['Condition'].isin(['earth', 'mars'])]
    data['morphology'] = df_morph[['Generation', 'Condition', 'Assay', 
                                    'length_50th', 'area_50th', 'width_midbody_50th']]
    
    # Load swimming data (Figure 3)
    df_swim = pd.read_excel(filepath, sheet_name='Figure 3')
    # Use the 'Swim freq' column
    df_swim = df_swim[df_swim['Generation'].isin([2.0, 4.0, 6.0])]
    df_swim = df_swim[df_swim['Condition'].isin(['earth', 'mars'])]
    data['swimming'] = df_swim[['Generation', 'Condition', 'Assay', 'Swim freq']].copy()
    data['swimming'].columns = ['Generation', 'Condition', 'Assay', 'swimming_frequency']
    
    # Load chemotaxis data (Figure 4)
    df_chemo = pd.read_excel(filepath, sheet_name='Figure 4')
    df_chemo = df_chemo[df_chemo['Generation'].isin([2.0, 4.0, 6.0])]
    df_chemo = df_chemo[df_chemo['Condition'].isin(['earth', 'mars'])]
    data['chemotaxis'] = df_chemo[['Generation', 'Condition', 'Assay', 'Chemotaxis Index']].copy()
    data['chemotaxis'].columns = ['Generation', 'Condition', 'Assay', 'chemotaxis_index']
    
    print(f"  Morphology: {len(data['morphology'])} observations")
    print(f"  Swimming: {len(data['swimming'])} observations")
    print(f"  Chemotaxis: {len(data['chemotaxis'])} observations")
    
    return data

def calculate_statistics(earth_data, mars_data):
    """
    Calculate t-test, effect size, and descriptive statistics.
    
    Parameters:
        earth_data (array): Data for Earth condition
        mars_data (array): Data for Mars condition
        
    Returns:
        dict: Dictionary with statistical results
    """
    # Descriptive statistics
    earth_mean = np.mean(earth_data)
    earth_std = np.std(earth_data, ddof=1)
    earth_sem = stats.sem(earth_data)
    earth_n = len(earth_data)
    
    mars_mean = np.mean(mars_data)
    mars_std = np.std(mars_data, ddof=1)
    mars_sem = stats.sem(mars_data)
    mars_n = len(mars_data)
    
    # T-test (two-tailed, independent samples)
    t_stat, p_value = stats.ttest_ind(earth_data, mars_data)
    df = earth_n + mars_n - 2
    
    # Cohen's d (pooled standard deviation)
    pooled_std = np.sqrt(((earth_n - 1) * earth_std**2 + (mars_n - 1) * mars_std**2) / df)
    cohens_d = (earth_mean - mars_mean) / pooled_std
    
    return {
        'earth_mean': earth_mean,
        'earth_std': earth_std,
        'earth_sem': earth_sem,
        'earth_n': earth_n,
        'mars_mean': mars_mean,
        'mars_std': mars_std,
        'mars_sem': mars_sem,
        'mars_n': mars_n,
        't_statistic': t_stat,
        'df': df,
        'p_value': p_value,
        'cohens_d': cohens_d,
        'significant': p_value < ALPHA
    }

def analyze_trait(data, trait_name, generations=[2, 4, 6]):
    """
    Analyze a single trait across generations.
    
    Parameters:
        data (DataFrame): Data with columns Generation, Condition, trait_name
        trait_name (str): Name of the trait column
        generations (list): Generations to analyze
        
    Returns:
        dict: Results for each generation
    """
    results = {}
    
    for gen in generations:
        gen_data = data[data['Generation'] == float(gen)]
        earth_data = gen_data[gen_data['Condition'] == 'earth'][trait_name].values
        mars_data = gen_data[gen_data['Condition'] == 'mars'][trait_name].values
        
        # Remove NaN values
        earth_data = earth_data[~np.isnan(earth_data)]
        mars_data = mars_data[~np.isnan(mars_data)]
        
        if len(earth_data) > 0 and len(mars_data) > 0:
            results[f'Gen{int(gen)}'] = calculate_statistics(earth_data, mars_data)
        else:
            print(f"Warning: No data for {trait_name} at Generation {gen}")
            
    return results

def create_verification_workbook(data, output_file):
    """
    Create Excel verification workbook with formulas.
    
    Parameters:
        data (dict): Dictionary with morphology, swimming, chemotaxis data
        output_file (str): Path to output Excel file
    """
    print(f"\nCreating verification workbook: {output_file}")
    
    with pd.ExcelWriter(output_file, engine='xlsxwriter') as writer:
        workbook = writer.book
        
        # Define formats
        header_format = workbook.add_format({
            'bold': True,
            'bg_color': '#4472C4',
            'font_color': 'white',
            'align': 'center',
            'border': 1
        })
        
        number_format = workbook.add_format({'num_format': '0.000'})
        pvalue_format = workbook.add_format({'num_format': '0.000'})
        
        # Create Overview sheet
        create_overview_sheet(writer, workbook, header_format)
        
        # Create analysis sheets
        create_morphology_sheets(writer, workbook, data['morphology'], header_format, number_format, pvalue_format)
        create_swimming_sheet(writer, workbook, data['swimming'], header_format, number_format, pvalue_format)
        create_chemotaxis_sheet(writer, workbook, data['chemotaxis'], header_format, number_format, pvalue_format)
        
    print(f"✓ Verification workbook created successfully")

def create_overview_sheet(writer, workbook, header_format):
    """Create overview/index sheet."""
    worksheet = workbook.add_worksheet('Overview')
    
    overview_data = [
        ['Statistical Verification Workbook'],
        ['Born on Mars: C. elegans Transgenerational Study'],
        [''],
        ['This workbook contains all statistical calculations reported in the manuscript.'],
        ['All values are calculated using Excel formulas for complete transparency.'],
        [''],
        ['CONTENTS:'],
        [''],
        ['Sheet', 'Description', 'Manuscript Reference'],
        ['Fig2A_Length', 'Body length (50th percentile) Earth vs Mars', 'Figure 2A'],
        ['Fig2B_Area', 'Body area (50th percentile) Earth vs Mars', 'Figure 2B'],
        ['Fig2C_Width', 'Midbody width (50th percentile) Earth vs Mars', 'Figure 2C'],
        ['Fig3_Swimming', 'Swimming frequency Earth vs Mars', 'Figure 3'],
        ['Fig4_Chemotaxis', 'Chemotaxis index Earth vs Mars', 'Figure 4'],
        [''],
        ['STATISTICAL METHODS:'],
        ['- Two-tailed independent t-tests'],
        ['- Cohen\'s d effect size (pooled standard deviation)'],
        ['- Significance level: α = 0.05'],
        ['- All formulas visible in cells'],
        [''],
        ['DATA SOURCE:'],
        ['Supplementary Data 1.xlsx'],
    ]
    
    for row_num, row_data in enumerate(overview_data):
        for col_num, value in enumerate(row_data):
            if row_num == 0:
                cell_format = workbook.add_format({'bold': True, 'font_size': 14})
                worksheet.write(row_num, col_num, value, cell_format)
            elif row_num == 8:
                worksheet.write(row_num, col_num, value, header_format)
            else:
                worksheet.write(row_num, col_num, value)
    
    worksheet.set_column('A:A', 20)
    worksheet.set_column('B:B', 50)
    worksheet.set_column('C:C', 20)

def create_morphology_sheets(writer, workbook, data, header_format, number_format, pvalue_format):
    """Create sheets for morphology traits."""
    
    traits = [
        ('Fig2A_Length', 'length_50th', 'Body Length (50th percentile)', 'μm'),
        ('Fig2B_Area', 'area_50th', 'Body Area (50th percentile)', 'μm²'),
        ('Fig2C_Width', 'width_midbody_50th', 'Midbody Width (50th percentile)', 'μm')
    ]
    
    for sheet_name, trait_col, trait_label, units in traits:
        worksheet = workbook.add_worksheet(sheet_name)
        
        # Title
        title_format = workbook.add_format({'bold': True, 'font_size': 12})
        worksheet.write(0, 0, trait_label, title_format)
        worksheet.write(1, 0, f'Units: {units}')
        
        generations = [2, 4, 6]
        
        for gen_idx, gen in enumerate(generations):
            start_row = 4 + gen_idx * 20
            
            # Section header
            gen_header_format = workbook.add_format({
                'bold': True,
                'bg_color': '#C5D9F1',
                'border': 1
            })
            worksheet.write(start_row, 0, f'GENERATION {int(gen)}', gen_header_format)
            
            # Extract data for this generation
            gen_data = data[data['Generation'] == float(gen)]
            earth_data = gen_data[gen_data['Condition'] == 'earth'][trait_col].dropna().values
            mars_data = gen_data[gen_data['Condition'] == 'mars'][trait_col].dropna().values
            
            # Data table
            row = start_row + 2
            worksheet.write(row, 0, 'Condition', header_format)
            for i in range(max(len(earth_data), len(mars_data))):
                worksheet.write(row, i+1, f'Replicate {i+1}', header_format)
            worksheet.write(row, len(earth_data)+2, 'Mean', header_format)
            worksheet.write(row, len(earth_data)+3, 'SD', header_format)
            worksheet.write(row, len(earth_data)+4, 'SEM', header_format)
            worksheet.write(row, len(earth_data)+5, 'N', header_format)
            
            # Earth data
            row += 1
            worksheet.write(row, 0, 'Earth')
            for i, val in enumerate(earth_data):
                worksheet.write(row, i+1, val, number_format)
            
            # Earth statistics with formulas
            earth_row_letter = row + 1
            worksheet.write_formula(row, len(earth_data)+2, 
                                   f'=AVERAGE(B{earth_row_letter}:F{earth_row_letter})', number_format)
            worksheet.write_formula(row, len(earth_data)+3, 
                                   f'=STDEV(B{earth_row_letter}:F{earth_row_letter})', number_format)
            worksheet.write_formula(row, len(earth_data)+4, 
                                   f'=I{earth_row_letter}/SQRT(K{earth_row_letter})', number_format)
            worksheet.write_formula(row, len(earth_data)+5, 
                                   f'=COUNT(B{earth_row_letter}:F{earth_row_letter})')
            
            # Mars data
            row += 1
            worksheet.write(row, 0, 'Mars')
            for i, val in enumerate(mars_data):
                worksheet.write(row, i+1, val, number_format)
            
            # Mars statistics with formulas
            mars_row_letter = row + 1
            worksheet.write_formula(row, len(mars_data)+2, 
                                   f'=AVERAGE(B{mars_row_letter}:F{mars_row_letter})', number_format)
            worksheet.write_formula(row, len(mars_data)+3, 
                                   f'=STDEV(B{mars_row_letter}:F{mars_row_letter})', number_format)
            worksheet.write_formula(row, len(mars_data)+4, 
                                   f'=I{mars_row_letter}/SQRT(K{mars_row_letter})', number_format)
            worksheet.write_formula(row, len(mars_data)+5, 
                                   f'=COUNT(B{mars_row_letter}:F{mars_row_letter})')
            
            # Statistical tests
            row += 2
            worksheet.write(row, 0, 'STATISTICAL TESTS', header_format)
            
            row += 1
            worksheet.write(row, 0, 'Degrees of freedom:')
            worksheet.write_formula(row, 1, f'=K{earth_row_letter}+K{mars_row_letter}-2')
            
            row += 1
            worksheet.write(row, 0, 'p-value:')
            worksheet.write_formula(row, 1, 
                                   f'=T.TEST(B{earth_row_letter}:F{earth_row_letter},B{mars_row_letter}:F{mars_row_letter},2,2)', 
                                   pvalue_format)
            
            row += 1
            worksheet.write(row, 0, 'Cohen\'s d:')
            # Cohen's d = (mean1 - mean2) / pooled_sd
            # pooled_sd = SQRT(((n1-1)*sd1^2 + (n2-1)*sd2^2) / (n1+n2-2))
            worksheet.write_formula(row, 1, 
                                   f'=(H{earth_row_letter}-H{mars_row_letter})/SQRT(((K{earth_row_letter}-1)*I{earth_row_letter}^2+(K{mars_row_letter}-1)*I{mars_row_letter}^2)/(K{earth_row_letter}+K{mars_row_letter}-2))', 
                                   number_format)
            
            row += 1
            worksheet.write(row, 0, 'Significant (α=0.05):')
            # Reference the p-value cell which is 2 rows up
            worksheet.write_formula(row, 1, f'=IF(B{row-2}<0.05,"YES","NO")')
        
        worksheet.set_column('A:A', 20)
        worksheet.set_column('B:I', 12)

def create_swimming_sheet(writer, workbook, data, header_format, number_format, pvalue_format):
    """Create sheet for swimming frequency."""
    create_behavioral_sheet(writer, workbook, data, 'swimming_frequency', 
                           'Fig3_Swimming', 'Swimming Frequency', 'Hz',
                           header_format, number_format, pvalue_format)

def create_chemotaxis_sheet(writer, workbook, data, header_format, number_format, pvalue_format):
    """Create sheet for chemotaxis index."""
    create_behavioral_sheet(writer, workbook, data, 'chemotaxis_index', 
                           'Fig4_Chemotaxis', 'Chemotaxis Index', 'index (0-1)',
                           header_format, number_format, pvalue_format)

def create_behavioral_sheet(writer, workbook, data, trait_col, sheet_name, trait_label, units,
                            header_format, number_format, pvalue_format):
    """Generic function to create behavioral trait sheets."""
    
    worksheet = workbook.add_worksheet(sheet_name)
    
    # Title
    title_format = workbook.add_format({'bold': True, 'font_size': 12})
    worksheet.write(0, 0, trait_label, title_format)
    worksheet.write(1, 0, f'Units: {units}')
    
    generations = [2, 4, 6]
    
    for gen_idx, gen in enumerate(generations):
        start_row = 4 + gen_idx * 20
        
        # Section header
        gen_header_format = workbook.add_format({
            'bold': True,
            'bg_color': '#C5D9F1',
            'border': 1
        })
        worksheet.write(start_row, 0, f'GENERATION {int(gen)}', gen_header_format)
        
        # Extract data for this generation
        gen_data = data[data['Generation'] == float(gen)]
        earth_data = gen_data[gen_data['Condition'] == 'earth'][trait_col].dropna().values
        mars_data = gen_data[gen_data['Condition'] == 'mars'][trait_col].dropna().values
        
        # Data table
        row = start_row + 2
        worksheet.write(row, 0, 'Condition', header_format)
        for i in range(max(len(earth_data), len(mars_data))):
            worksheet.write(row, i+1, f'Replicate {i+1}', header_format)
        worksheet.write(row, len(earth_data)+2, 'Mean', header_format)
        worksheet.write(row, len(earth_data)+3, 'SD', header_format)
        worksheet.write(row, len(earth_data)+4, 'SEM', header_format)
        worksheet.write(row, len(earth_data)+5, 'N', header_format)
        
        # Earth data
        row += 1
        worksheet.write(row, 0, 'Earth')
        for i, val in enumerate(earth_data):
            worksheet.write(row, i+1, val, number_format)
        
        # Earth statistics
        earth_row_letter = row + 1
        worksheet.write_formula(row, len(earth_data)+2, 
                               f'=AVERAGE(B{earth_row_letter}:F{earth_row_letter})', number_format)
        worksheet.write_formula(row, len(earth_data)+3, 
                               f'=STDEV(B{earth_row_letter}:F{earth_row_letter})', number_format)
        worksheet.write_formula(row, len(earth_data)+4, 
                               f'=I{earth_row_letter}/SQRT(K{earth_row_letter})', number_format)
        worksheet.write_formula(row, len(earth_data)+5, 
                               f'=COUNT(B{earth_row_letter}:F{earth_row_letter})')
        
        # Mars data
        row += 1
        worksheet.write(row, 0, 'Mars')
        for i, val in enumerate(mars_data):
            worksheet.write(row, i+1, val, number_format)
        
        # Mars statistics
        mars_row_letter = row + 1
        worksheet.write_formula(row, len(mars_data)+2, 
                               f'=AVERAGE(B{mars_row_letter}:F{mars_row_letter})', number_format)
        worksheet.write_formula(row, len(mars_data)+3, 
                               f'=STDEV(B{mars_row_letter}:F{mars_row_letter})', number_format)
        worksheet.write_formula(row, len(mars_data)+4, 
                               f'=I{mars_row_letter}/SQRT(K{mars_row_letter})', number_format)
        worksheet.write_formula(row, len(mars_data)+5, 
                               f'=COUNT(B{mars_row_letter}:F{mars_row_letter})')
        
        # Statistical tests
        row += 2
        worksheet.write(row, 0, 'STATISTICAL TESTS', header_format)
        
        row += 1
        worksheet.write(row, 0, 'Degrees of freedom:')
        worksheet.write_formula(row, 1, f'=K{earth_row_letter}+K{mars_row_letter}-2')
        
        row += 1
        worksheet.write(row, 0, 'p-value:')
        worksheet.write_formula(row, 1, 
                               f'=T.TEST(B{earth_row_letter}:F{earth_row_letter},B{mars_row_letter}:F{mars_row_letter},2,2)', 
                               pvalue_format)
        
        row += 1
        worksheet.write(row, 0, 'Cohen\'s d:')
        worksheet.write_formula(row, 1, 
                               f'=(H{earth_row_letter}-H{mars_row_letter})/SQRT(((K{earth_row_letter}-1)*I{earth_row_letter}^2+(K{mars_row_letter}-1)*I{mars_row_letter}^2)/(K{earth_row_letter}+K{mars_row_letter}-2))', 
                               number_format)
        
        row += 1
        worksheet.write(row, 0, 'Significant (α=0.05):')
        worksheet.write_formula(row, 1, f'=IF(B{row-2}<0.05,"YES","NO")')
    
    worksheet.set_column('A:A', 20)
    worksheet.set_column('B:I', 12)

def print_results_summary(data):
    """Print comprehensive summary of all statistical results."""
    
    print("\n" + "="*80)
    print("STATISTICAL RESULTS SUMMARY")
    print("="*80)
    
    # Morphology
    print("\n### MORPHOLOGY (Figure 2) ###\n")
    
    traits = [
        ('length_50th', 'Body Length'),
        ('area_50th', 'Body Area'),
        ('width_midbody_50th', 'Midbody Width')
    ]
    
    for trait_col, trait_label in traits:
        print(f"\n{trait_label}:")
        print("-" * 60)
        results = analyze_trait(data['morphology'], trait_col)
        
        for gen_key, stats in results.items():
            sig_marker = "***" if stats['p_value'] < 0.001 else \
                        "**" if stats['p_value'] < 0.01 else \
                        "*" if stats['p_value'] < 0.05 else "ns"
            
            print(f"\n{gen_key}:")
            print(f"  Earth: {stats['earth_mean']:.2f} ± {stats['earth_sem']:.2f} (n={stats['earth_n']})")
            print(f"  Mars:  {stats['mars_mean']:.2f} ± {stats['mars_sem']:.2f} (n={stats['mars_n']})")
            print(f"  t({stats['df']}) = {stats['t_statistic']:.3f}, p = {stats['p_value']:.4f} {sig_marker}")
            print(f"  Cohen's d = {stats['cohens_d']:.2f}")
    
    # Swimming
    print("\n\n### SWIMMING FREQUENCY (Figure 3) ###\n")
    print("-" * 60)
    results = analyze_trait(data['swimming'], 'swimming_frequency')
    
    for gen_key, stats in results.items():
        sig_marker = "***" if stats['p_value'] < 0.001 else \
                    "**" if stats['p_value'] < 0.01 else \
                    "*" if stats['p_value'] < 0.05 else "ns"
        
        print(f"\n{gen_key}:")
        print(f"  Earth: {stats['earth_mean']:.3f} ± {stats['earth_sem']:.3f} Hz (n={stats['earth_n']})")
        print(f"  Mars:  {stats['mars_mean']:.3f} ± {stats['mars_sem']:.3f} Hz (n={stats['mars_n']})")
        print(f"  t({stats['df']}) = {stats['t_statistic']:.3f}, p = {stats['p_value']:.4f} {sig_marker}")
        print(f"  Cohen's d = {stats['cohens_d']:.2f}")
    
    # Chemotaxis
    print("\n\n### CHEMOTAXIS INDEX (Figure 4) ###\n")
    print("-" * 60)
    results = analyze_trait(data['chemotaxis'], 'chemotaxis_index')
    
    for gen_key, stats in results.items():
        sig_marker = "***" if stats['p_value'] < 0.001 else \
                    "**" if stats['p_value'] < 0.01 else \
                    "*" if stats['p_value'] < 0.05 else "ns"
        
        print(f"\n{gen_key}:")
        print(f"  Earth: {stats['earth_mean']:.3f} ± {stats['earth_sem']:.3f} (n={stats['earth_n']})")
        print(f"  Mars:  {stats['mars_mean']:.3f} ± {stats['mars_sem']:.3f} (n={stats['mars_n']})")
        print(f"  t({stats['df']}) = {stats['t_statistic']:.3f}, p = {stats['p_value']:.4f} {sig_marker}")
        print(f"  Cohen's d = {stats['cohens_d']:.2f}")
    
    print("\n" + "="*80)
    print("Analysis complete!")
    print("="*80 + "\n")

def main():
    """Main analysis function."""
    
    print("\n" + "="*80)
    print("MARS WORMS STATISTICAL ANALYSIS")
    print("="*80)
    
    # Check if input file exists
    if not Path(INPUT_FILE).exists():
        print(f"\nERROR: Input file '{INPUT_FILE}' not found!")
        print("Please ensure Supplementary Data 1.xlsx is in the current directory.")
        sys.exit(1)
    
    # Load data
    data = load_data(INPUT_FILE)
    
    # Print results
    print_results_summary(data)
    
    # Create verification workbook
    create_verification_workbook(data, OUTPUT_FILE)
    
    print(f"\n✓ All analyses complete!")
    print(f"✓ Verification workbook saved: {OUTPUT_FILE}")
    print(f"\nYou can now submit both files as supplementary materials:\n")
    print(f"  Supplementary Data 1: {INPUT_FILE}")
    print(f"  Supplementary Data 2: {OUTPUT_FILE}")
    print(f"  Supplementary Data 3: {Path(__file__).name}")

if __name__ == "__main__":
    main()
```
